# Metacommunity-scale biodiversity regulation and the self-organized emergence of macroecological patterns

**DOI:** 10.1101/489336

**Authors:** Jacob D. O’Sullivan, Robert J. Knell, Axel G. Rossberg

## Abstract

There exist a number of key macroecological patterns whose ubiquity suggests the spatio-temporal structure of ecological communities is governed by some universal mechanisms. The nature of these mechanisms, however, remains poorly understood. Here we probe spatio-temporal patterns in species richness and community composition using a simple metacommunity assembly model. Despite making no *a priori* assumptions regarding biotic spatial structure or the distribution of biomass across species, model metacommunities self-organize to reproduce well documented patterns including characteristic species abundance distributions, range size distributions and species area relations. Also in agreement with observations, species richness in our model attains an equilibrium despite continuous species turnover. Crucially, it is in the neighbourhood of the equilibrium that we observe the emergence of these key macroecological patterns. Biodiversity equilibria in models occur due to the onset of ecological structural instability, a population-dynamical mechanism. This strongly suggests a causal link between local community processes and macroecological phenomena.

Should this manuscript be accepted all simulation data supporting the results will be archived in a public repository and the data DOI will be included at the end of the article

## Introduction

Despite the colossal diversity of environments where life is found across the globe, there exist a number of spatio-temporal patterns in biodiversity which are observed in almost every ecological community that has been studied. The species abundance distribution (SAD), which highlights the overwhelming predominance of rare species in ecological communities, has long been considered universal (Fisher et al., 1943; Preston, 1948). A related pattern, the range size distribution (RSD), points to the prevalence of small (Brown et al., 1996; Gaston, 1996), aggregated (Brown, 1984) ranges, with few species occupying broad distributions. The species area relation (SAR), which denotes the sub-linear increase in diversity as a function of sample area, has been described as “one of community ecology’s few genuine laws” (Schoener, 1976).

The less extensively studied (and considerably more divisive) phenomenon of community-level diversity regulation, which constrains the number of species coexisting within an assemblage, has recently been proposed as a general ecological pattern (Gotelli et al., 2017; Magurran et al., 2018). Evidence of strong diversity regulation has been found in desert rodents (Brown et al., 2000), birds (Parody et al., 2001), marine fish (Magurran et al., 2015), freshwater communities (Magurran et al., 2018), and in global-scale meta-analyses (Dornelas et al., 2014; Gotelli et al., 2017). At geological timescales, constrained diversification, assumed to reflect the impact of ecological limits on evolutionary processes, has been detected in the fossil record of a variety of taxa (Alroy, 2009, 2010; Liow and Finarelli, 2014; Benson et al., 2016; Close et al., 2019). Despite their apparent ubiquity, which strongly hints at some almost universal processes in macroecology, our mechanistic understanding of these spatio-temporal patterns in biodiversity remains disparate and incomplete.

The theory of island biogeography (MacArthur and Wilson, 1967) suggests that patterns in biodiversity may be explained as a consequence of the dependence of diversification rates - speciation, invasion and extinction - on standing diversity. In the fifty years since the Theory of Island Biogeography was first developed, however, the effects of environmental heterogeneity, landscape topography, local species interactions and dispersal have been shown to impact local and regional diversity patterns in complex ways (Shmida and Wilson, 1985; Holt, 1985; Pulliam, 1988). Contemporary metacommunity ecology (Leibold et al., 2004; Holyoak et al., 2005; Logue et al., 2011; Winegardner et al., 2012) shines a light on how these complex and overlapping processes interact. Perhaps due to the persistent view that local and regional ecological processes cannot be meaningfully unified (Harmon and Harrison, 2015), surprisingly few studies consider metacommunity frameworks that explicitly incorporate community dynamics at multiple spatial scales (e.g. Pillai et al., 2010; Barter and Gross, 2017; but see Plitzko and Drossel, 2015; Thiel and Drossel, 2018). Here we attempt to fill this gap with a dynamically simple metacommunity assembly model which incorporates local ecological interactions and dispersal in an environmentally heterogeneous landscape, thus uniting the branches of population-dynamical and spatial ecology. Our primary focus in developing this model was the study of how biodiversity might be regulated in spatially resolved ecological assemblages, but we find an intriguing emergent relationship between diversity regulation in model metacommunities and the appearance of widely observed macroecological patterns.

In spatially *unresolved* models, the emergence of biodiversity regulation has been observed numerous times (e.g. Drossel et al., 2001; Yoshida, 2003; Pawar, 2009). Analytic theory (Rossberg, 2013) reveals that this phenomenon is caused by the loss of *ecological structural stability*, which denotes the robustness of assemblages to press (i.e. sustained) perturbations (Meszéna et al., 2006; Bastolla et al., 2005, 2009; Rossberg, 2013; Rohr et al., 2014; Barbier et al., 2018). The mechanism is most easily understood for the paradigmatic case of Lotka-Volterra competition models of the form

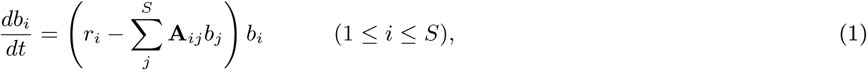

with population biomasses *b*_*i*_, linear growth rates *r*_*i*_, competition coefficients **A**_*ij*_ *≥* 0, and species richness *S*. If all *S* species co-exist (*b*_*i*_ *>* 0, for all *i*), the equilibrium condition for this system can be written in matrix-vector notation as **r** - **Ab** = 0 and is solved by **b** = **A**^-1^**r**. Mathematical problems of this form are called ‘ill conditioned’, implying that the solution **b** responds sensitively to changes in both **A** and **r**, when some eigenvalues of **A** approach zero. In ecological models we define this sensitivity as ecological structural *in*stability. Once this unstable condition arises, perturbation by external pressures or invaders, formally presentable by changes in **A** or **r**, can easily lead to extinctions. Structural stability (controlled by **A**) and linear/Lyapunov stability (controlled by the Jacobian matrix) should not be confused. While these two phenomena are related (Stone, 2018), each of them can independently control community structure and dynamics.

Random matrix theory of the kind invoked by May (1973), but applied to the competition matrix **A** rather than the system’s Jacobian matrix at equilibrium, robustly predicts that with increasing species richness some eigenvalues of **A** approach zero. The overall effect is thus that with increasing species richness structural instability increases, and accordingly, the likelihood that invasions cause species extinctions. Community assembly models therefore converge on dynamic steady states defined by the onset of structural instability. By applying alternative mathematical approaches to studying this phenomenon (Yodzis, 1988; Tokita, 2004; Rossberg, 2013; Dougoud et al., 2018; Barbier et al., 2018; Galla, 2018), structural instability can be interpreted as resulting from the amplification of perturbations through complex indirect interactions in large communities.

For other types of spatially unresolved community models, approximation techniques have been developed to map these onto competition models of the form Eq. (1) (layered food webs: Bastolla et al. 2005, mutualistic communities: Bastolla et al. 2009, arbitrary food webs: Rossberg 2013), and the theory applies analogously. Whether this is similarly the case for metacommunity models is unknown. In order to understand the relationship between diversity regulation, potentially via the onset of ecological structural instability, and processes active in metacommunities, we constructed a multi-species framework in which metapopulation dynamics at the regional scale are modelled using a spatial network of Lotka-Volterra competition equations with additional terms describing dispersal (see *Dynamic equations and metacommunity assembly*, below).

By comparing the model’s behaviour to analytic predictions developed for spatially unresolved competitive communities (Rossberg, 2013), we show that intrinsic metacommunity-level diversity regulation can indeed be explained as a consequence of the onset of ecological structural instability at the *regional* scale. Surprisingly, and potentially very importantly, we find that, as model metacommunities approach diversity limits, they *self organize* to reproduce macroecological patterns previously identified as central for the spatial structure of biodiversity (McGill, 2010): a skewed local and regional distribution of abundances, spatial aggregation of conspecific biomass, and apparent absence of species co-occurrence patterns. In combination, as McGill (2010) argued, these core patterns lead to sub-linear species area relations and other spatial biodiversity phenomena. That a diverse set of well known macroecological patterns emerges in a simple metacommunity model strongly supports the hypothesis that these patterns are indeed indirect consequences of local, niche-based population dynamics and dispersal.

## Results and discussion

### Metacommunity species richness

Simulated metacommunities, assembled in our model via a constant, slow influx of invaders, converge on regional diversity equilibria at which species richness remains approximately stationary despite continuous turnover in composition (Fig. 1). Diversity relaxes back to the same approximate steady state after sudden removal or introduction of large numbers of species (Fig. 1).

**Figure 1:**
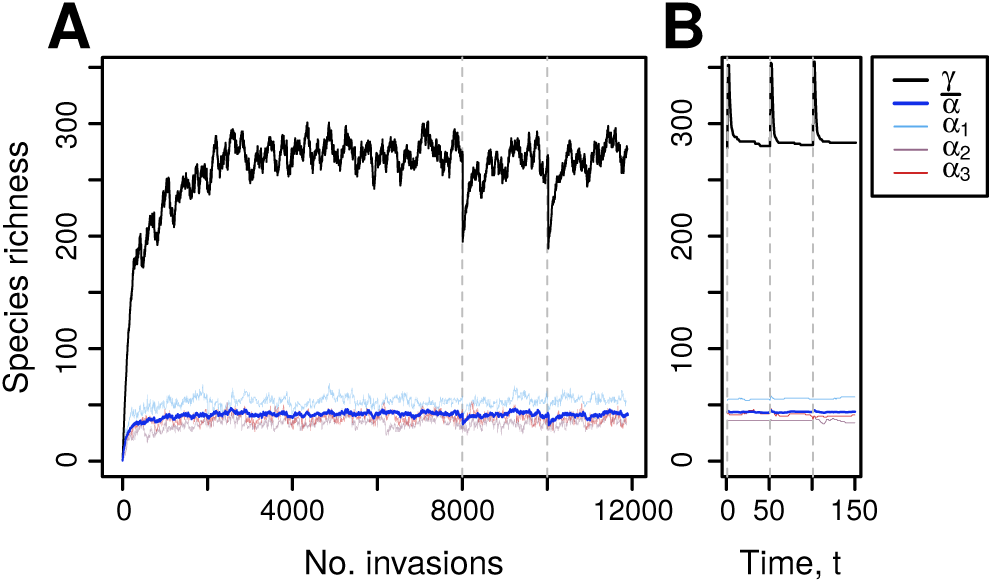
Biodiversity regulation in model metacommunities. The emergence of diversity equilibria at multiple spatial scales as a result of a stepwise invasion flux in a typical model metacommunity (A). Regional diversity (γ, black) and the average local diversity (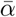, blue) are shown, as well as that observed in three randomly selected patches (α, coloured). Relaxation back to equilibrium following random removal or introduction of large numbers of species (25% of the equilibrium richness, indicated by vertical dashed lines, A and B) reveals the strength and predictability of metacommunity-scale regulation in model assemblages. Relaxation times following removal and introduction differ by several orders of magnitude since the re-accumulation of diversity occurs at the invasion timescale, while extirpations occur at the population-dynamic timescale (measured in units *t*).

From previous theoretical work we know that the sensitivity of a spatially unresolved community to press perturbations is a function of the standing diversity and the intensity of ecological interactions within that community (Rossberg, 2013). The structurally unstable limit around which diversity in model communities converges, denoted *S\**, is a function of the statistical distribution of the competition coefficients, typically its first and second moments (mean, variance, covariances). By assuming a precisely analogous mechanism to operate at the metacommunity scale, the basic spatially unresolved theory predicts an approximate *regional* diversity of

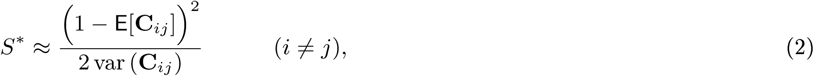

(Rossberg, 2013, Eq. 17.5) where E[**C**_*ij*_] and var (**C**_*ij*_) represent the expectation and variance of the interspecific competition coefficients computed at the scale of the metacommunity, replacing the distribution of local interaction coefficients **A**_*ij*_ in the spatially unresolved theory. The eigenvalue spectrum of the competitive overlap matrix offers a convenient graphical tool for assessing the ecological structural stability of community models. The structurally unstable diversity limit occurs as the area covered by the spectrum in the complex plane approach the origin (Rossberg, 2013).

Regional-scale, interspecific competition coefficients **C**_*ij*_ were computed for model metacommunities assembled for a range of parameter combinations, and used to evaluate Eq. (2) (see *Regional scale interaction matrices*; **C** below). Comparing the diversity predicted by Eq. (2) with that in the steady state of the simulation, we found that the spatially unresolved analytic prediction explains 95% of variance in the equilibrium species richness in the spatially resolved models (Fig. 2A). Furthermore, the spectra of the matrices **C** approach the origin when the biodiversity equilibrium is reached (Fig. 2B), just as observed in spatially unresolved models (Rossberg, 2013). Thus, although an analytic prediction of the structurally unstable diversity limit in the metacommunity case is not available due to the intractability of the full model, we find strong evidence supporting the claim that ecological structural stability drives diversity regulation at the metacommunity scale.

**Figure 2:**
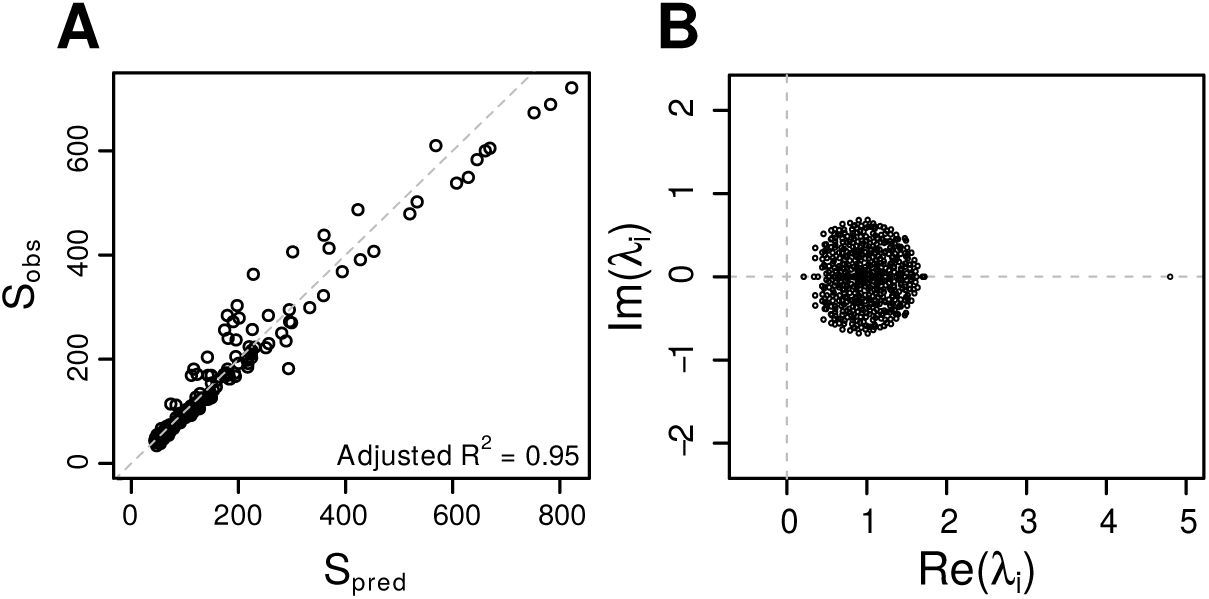
Testing for biodiversity regulation by structural stability in metacommunity models. **A**: Comparison of the regional equilibrium diversity predicted by Eq. (2) and that observed in simulated metacommunities for 195 combinations of patch number and spatial heterogeneity. The dashed line signifies equality. **B**: The eigenvalue spectrum of a typical regional-scale competitive overlap matrix **C**. Both analyses strongly suggest the mechanism regulating diversity at the metacommunity scale is the loss of ecological structural stability.

The relationship between local and regional competition coefficients is non-trivial and depends on the degree of environmental heterogeneity (see *Model landscape*, below). Nonetheless, we found the off-diagonal elements of **A**_*ij*_ and **C**_*ij*_ to be significantly correlated in metacommunity models at regional diversity limits (*p <* 0.01, for all parameter combinations). This implies that local ecological interactions propagate to the metacommunity scale and influence regional diversity patterns (Rabosky and Hurlbert, 2015). For further discussion see Supporting Information.

### Local species richness

In order to distinguish between local and regional diversity it is necessary to define some criterion for assessing presence-absence in a local assemblage. We do this in two ways. First by setting an arbitrary limit, equivalent to a detection threshold, of 10^-4^ biomass units below which a species is considered to be absent from a local community. This value is four orders of magnitude lower than the maximum local biomass permitted in the model and therefore defines a detectable range that is in accordance with many empirical observations (e.g. Condit et al., 2002). We further distinguish among those populations exceeding the detection threshold by defining *source* and *sink* populations as those capable of self maintenance in a given location, and those that would decline without continuous immigration from adjacent communities (see *Source-Sink classification*, below).

During the assembly process (Fig. 1), we find that local community richness, defined by the detection threshold, saturates earlier than the regional assemblage (after around ∼ 500 and ∼ 4000 invasions, respectively, in the example shown). To confirm whether this local community regulation occurs independently of metacommunity regulation, it is necessary to ask how *α*-diversity is related to *γ*-diversity. If local diversity limits are controlled indirectly by the size of the regional species pool (i.e. regulation occurs meaningfully at one scale only) we would expect a linear, or at least non-saturating local-regional species richness relation. Interestingly, however, both source and sink diversity, and, by extension, their sum, are saturating functions of the regional species richness in equilibrial metacommunities. As we show in Fig. 3A, for which the distribution of interspecific coefficients **A**_*ij*_ was fixed across all simulations and considering source populations only, average local diversity converged on a horizontal asymptote of ∼ 50 once the regional assemblage reached ∼ 300 species. Similar convergence, though with greater scatter, is evident for sink populations, though the asymptote may occur outside of the range studied here. This suggests that local diversity is indeed independently regulated, such that in sufficiently large regional communities local diversity is effectively independent of the metacommunity-scale parameterization that determines the size of the regional species pool.

**Figure 3:**
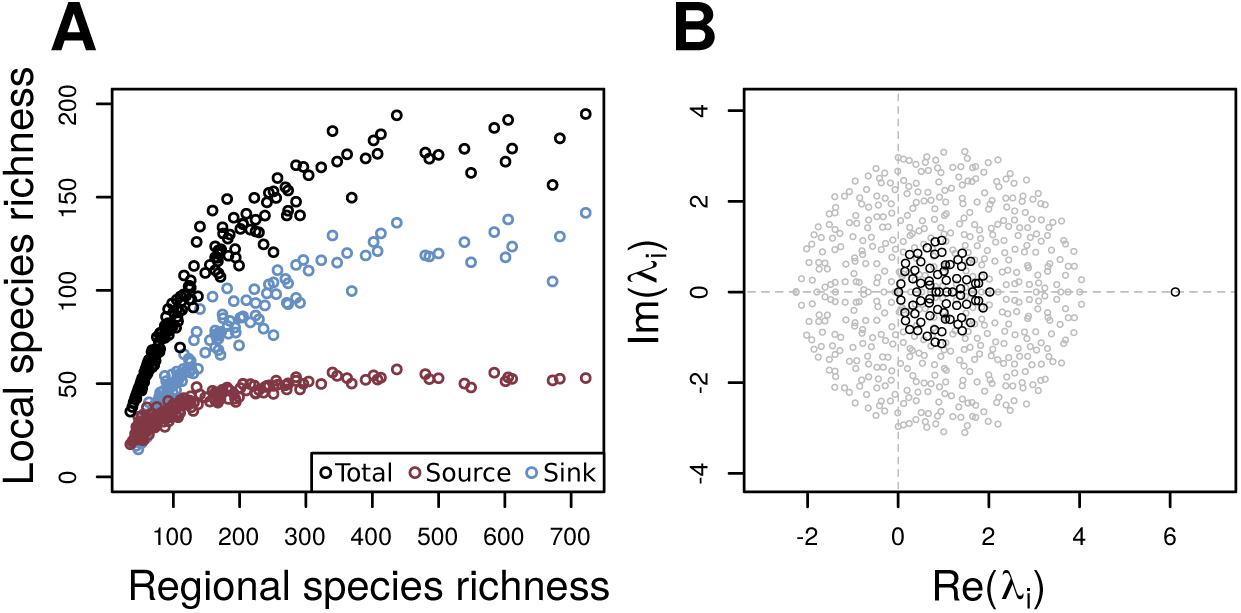
Demonstration of structurally unstable diversity regulation at the local scale. A: Average local diversity (black), and that attributed to source (red) and sink (blue) populations, at regional diversity equilibrium, plotted against regional species richness for the same 195 parameter combinations used in Fig. 2. The sublinearity of the local-regional richness relation suggests local communities are saturated with respect to both source and sink diversity for sufficiently high *N*. B: Comparison of the spectra of the full competitive overlap matrix **A** (grey circles) with that of its sub-matrix matrix **A**_source_ (black circles) corresponding to source populations only, for a randomly selected local community at regional diversity equilibrium. The spectrum of **A**_source_ demonstrates the role of structural instability in regulating the diversity of the key, locally sustained component of the local assemblage.

To explore the mechanism responsible for local diversity regulation we determined, for any given patch, the sub-matrix of **A** corresponding to the local source populations only. We found that the spectra of these sub-matrices, too, approach the origin of the complex plane. Thus we concluded that structurally unstable dynamics regulate species richness not only at the metacommunity level, but also, independently at the local level (Fig. 3B), and define sink populations as the super-saturated component of the local assemblage which depends on non-equilibrium dynamics (mass effects) for persistence.

### Temporal turnover

Stationarity in species richness is a key and unambiguous characteristic of our model metacommunities. Less obvious is the fact that, rather than converging on a near-static, ‘climax’ community, metacommunity composition in our models continuously turns over in response to the slow flux of invaders (Fig. 4A). Interestingly, on average, local communities turn over faster than the regional metacommunity of which they form a part. This is seen in the rapid decay in community similarity at the local, relative to the regional scale (Fig. 4A). This might be explained by range contraction and expansion due to regional biotic turnover which occur faster than landscape-scale competitive exclusion.

**Figure 4:**
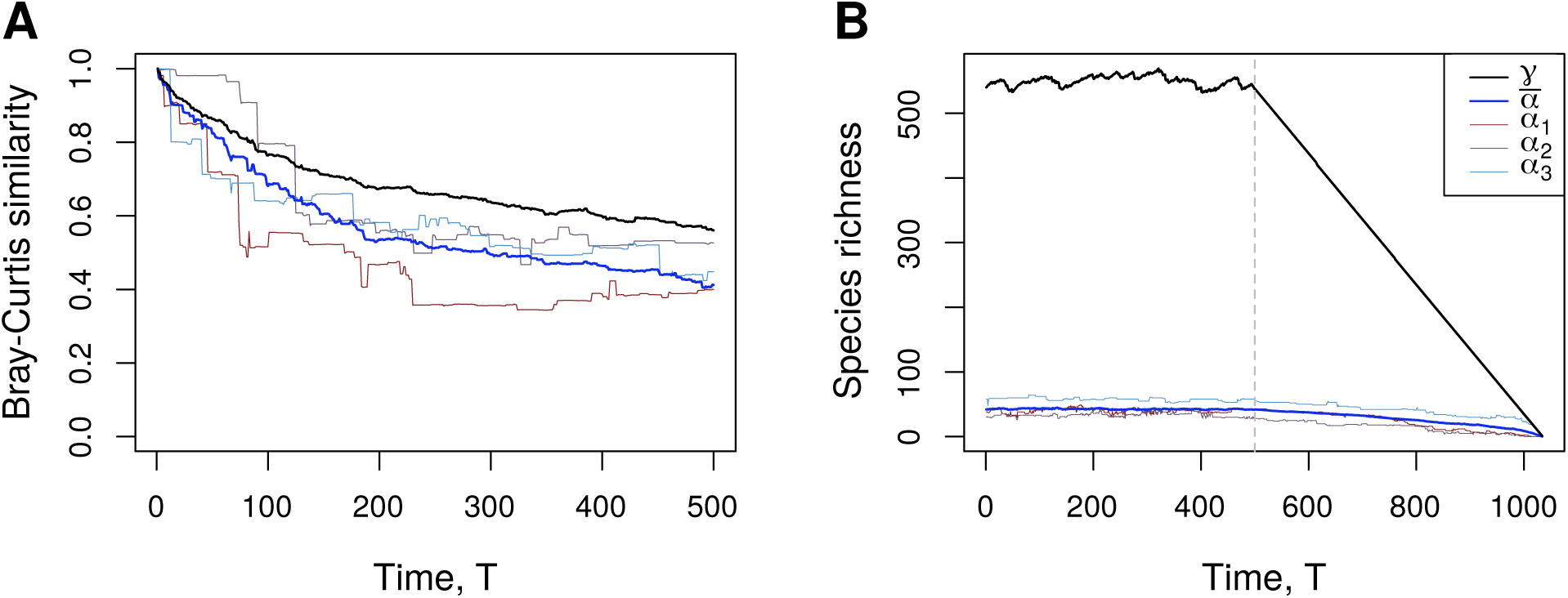
Temporal trends in community composition and species richness of model communities. The temporal Bray-Curtis similarity (A) and *source* population species richness (B, *T <* 500) for a metacommunity at regional equilibrium subject to a slow, discrete flux of invaders. Data for the metacommunity (black), the local average (blue) and three randomly selected local communities (coloured) are shown. After 500 invasions the invasion flux in panel (B) is switched off and species are successively removed in order of increasing regional abundance.

If the invader flux is spontaneously stopped and species are experimentally removed from the metacommunity in increasing order of regional biomass, fast turnover at local scales buffers local communities from the diversity losses at metacommunity level (Fig. 4B). In the example shown in Fig. 4, a 20% decrease in regional species richness produced only a 13% drop at the local scale on average. It has been estimated that the current rate of global species loss is 100-1000 times the background rate (De Vos et al., 2015; Pimm et al., 2014; Ceballos et al., 2015), yet global meta-analyses have failed to detect a consistent loss of diversity at the local scale (Dornelas et al., 2014; Vellend et al., 2017; Gotelli et al., 2017). Our results suggest that independent regulatory processes operating at multiple spatial scales may account for the discrepancy between local and regional/global diversity trends.

### Spatial patterns in biodiversity and abundance

By elegantly comparing the various major efforts to devise unified macroecological theory to date, McGill (2010) showed that three key macroecological phenomena are basic assumptions implicit to all frameworks. McGill argued that these key phenomena on their own are sufficient to give rise to a variety of emergent macroecological patterns, such as the sub-linear SAR. The three key patterns are an uneven SAD at various spatial scales, the spatial aggregation of conspecific biomass underlying the observed skewed RSD, and the (apparent) non-significant correlation in species’ spatial distributions. The last phenomenon relates to the observation that statistically significant positive or negative correlations in species’ spatial distributions or co-occurrence patterns are surprisingly under-represented in empirical studies; most pair-wise correlations tend to be indistinguishable from random (Hoagland and Collins, 1997; Veech, 2006; Houlahan et al., 2007; D’Amen et al., 2018). We find that, surprisingly, each of these three patterns emerges in our model metacommunities in the neighbourhood of regional diversity equilibria (Fig. 5).

**Figure 5:**
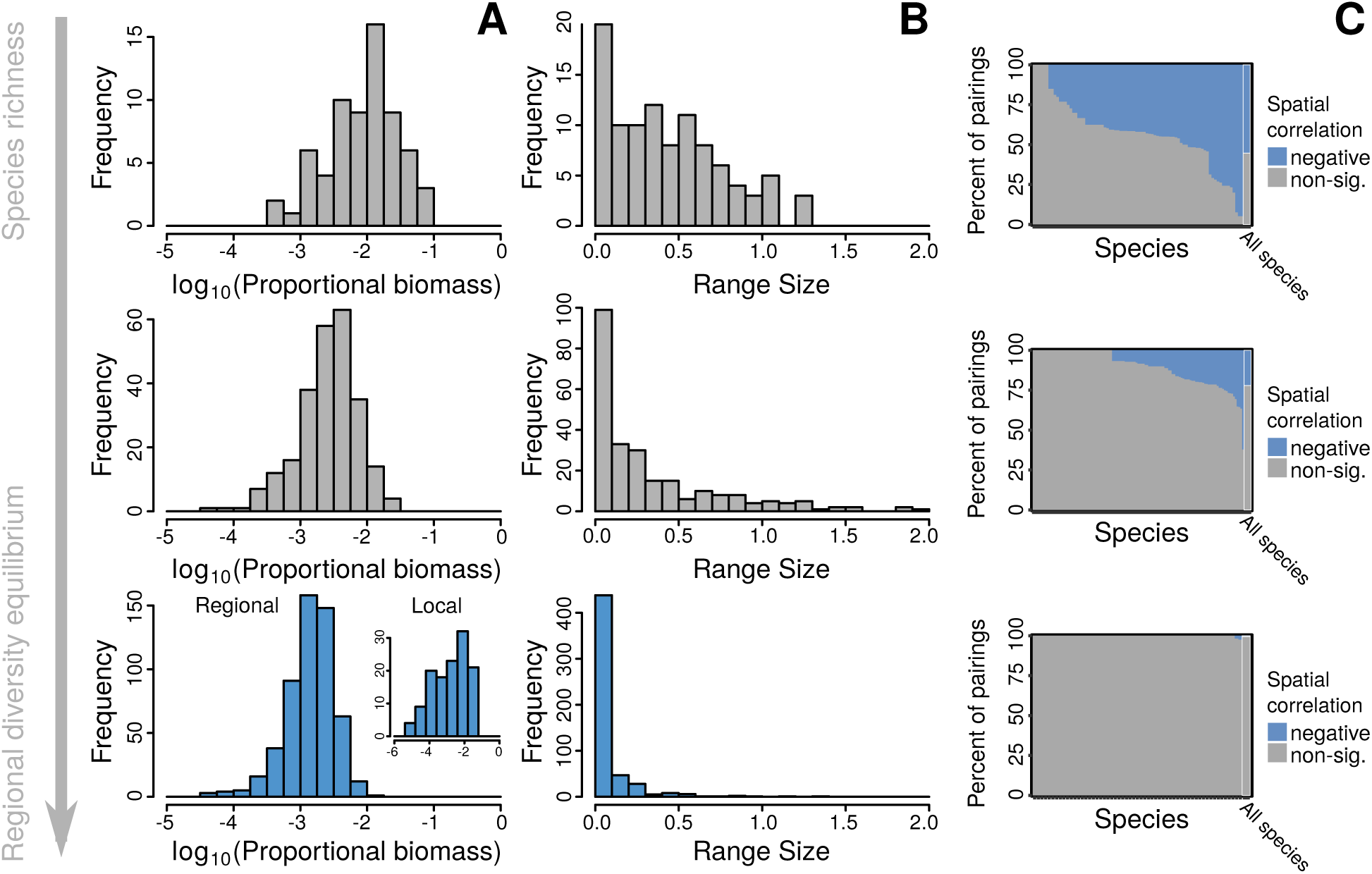
The effect of regional diversity regulation on macroecology in simulated metacommunities. Species Biomass Distributions (A), Range Size Distributions (B), and (C) Co-occurrence Profiles (Veech, 2006; Griffith et al., 2016) for a typical model metacommunity at 20, 50, and 100% of the regional diversity equilibrium of around 500 species. In B, an even distribution over the model landscape corresponds to a Range Size measure of 1.7, though for species concentrated near the edges our measure of Range Size can give even larger values. In C, the percentage of possible species pairs that exhibit statistically significant (*p <* 0.05) negative spatial correlation is shown in blue for each species, and for the community as a whole (right-most bar). Unsurprisingly, given the purely competitive nature of local ecological interactions, in our model metacommunties, no significant positive correlations were found. All three distributions converge on patterns well represented in the empirical literature as metacommunities approach the self-organized equilibrium.

Figure 5A shows that at both local and regional scale the SAD are left-skewed log-normal, as observed in communities ranging from marine benthos to Amazonian rain forest (McGill et al., 2007). The early onset of diversity regulation at the local scale already leads to highly skewed SAD at the regional scale, after which further accumulation of diversity at the regional scale drives the distribution to the left as average biomasses decline.

In the early stages of the assembly process, as local diversity accumulates, weak biotic filtering means species disperse across much of their fundamental geographic niche. Once local communities become constrained, regional invasions instead drive an increase in spatial *β*-diversity, the ‘regionalization’of the biota (Ricklefs, 2004), and a corresponding reduction of species ranges, which become highly spatially aggregated. At the metacommunity scale this is seen as a collapse in the RSD as the assemblage approaches regional diversity equilibrium (Fig. 5B). The skewed RSD for metacommunity models at regional diversity limits match patterns observed for a wide variety of taxa (Gaston, 1998), including pine species (Brown et al., 1996), tropical tree species (Xu et al., 2015) and in both regional (Gaston, 1996), and global distributions (Orme et al., 2006) of bird species. Reduction in average range sizes and the corresponding increase in the number of effective interactions with neighbouring populations may increase species’vulnerability to regional extinction (Ricklefs, 2004). As such we consider the emergent spatial aggregation in our metacommunity models to play an important role in regional scale diversity regulation.

The dependence of RSD on species richness implies a strong impact of ecological interactions on species ranges. Counter-intuitively, however, the vast majority of species pairs show no significant spatial correlation (Fig. 5C). As strong regional scale diversity regulation sets in and spatial ranges collapse, the percentage of species pairs for which it is possible to detect non-random spatial correlation drops to near zero, giving the impression of an eminently neutral system.

We considered the possibility that this absence of demonstrable spatial correlations is explained by the systemic exclusion of competing species pairs during assembly. However, for fully assembled model metacommunities at the regional diversity limit, non-zero interspecific competition coefficients made up 21.5-28.3% of the elements of the matrix **A**_*ij*_, only 1.7-8.5% less than in the statistical ensemble from which invaders are sampled.

McGill (2010) argues that these three key phenomena (Fig. 5) can combine to produce sub-linearity in the SAR. A recent non-dynamical modelling approach has verified that spatial aggregation at the population scale can indeed generate high level macroecological configurations (Takashina et al., 2019). Here we build on that result by showing that population-dynamical processes can drive spatial aggregation and produce the characteristic relationship between diversity and landscape area. The SAR in our models (Fig. 6) are well approximated by power laws with exponents ranging from 0.19 to 0.87, depending on the degree of spatial correlation in the model environment: a spatially more correlated, homogeneous environment produces an SAR with lower exponent. The exponents that emerge are well within the range found in a meta-analysis of almost 800 empirical SARs (Drakare et al., 2006). It has been shown (Rosindell and Cornell, 2007; Pigolotti et al., 2018) that realistic SAR emerge in spatially explicit neutral models. Here we show, in light of evidence against neutral community assembly at regional scales (Ostling, 2005), how sublinearity in the SAR can emerge via an explicit diversity dependent mechanism.

**Figure 6:**
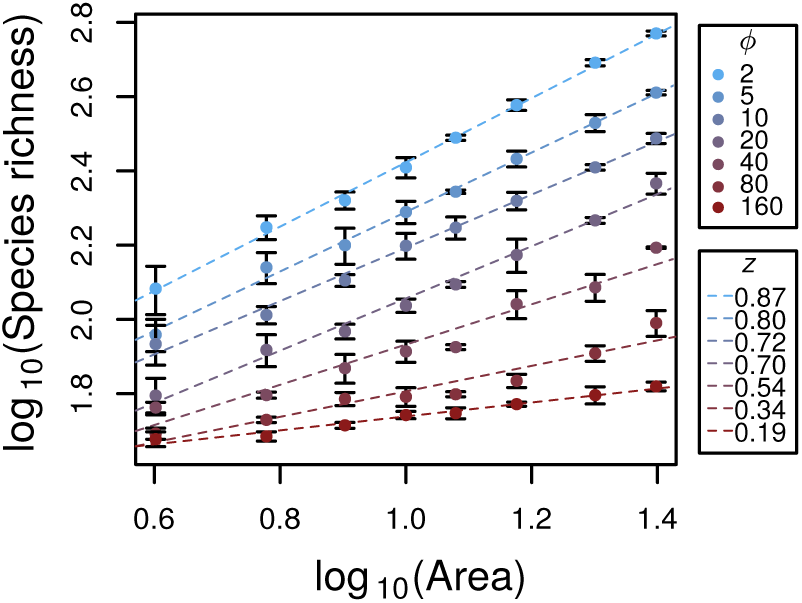
Species area relations for simulated metacommunities. Species richness increases as a function of model area according to approximate power laws with exponents ranging from 0.19 to 0.87. Colours indicate the degree of spatial environmental heterogeneity, *ϕ*. Error bars indicate standard deviations estimated from three independent model runs for each parameterization, for which small differences between simulations arose largely due to the random topography of the model landscapes.

## Conclusions

There is a growing body of evidence indicating that community level diversity regulation is a common characteristic of ecological communities at both local and regional scales (e.g. Alroy, 2009; Magurran et al., 2015, 2018; Gotelli et al., 2017; Dornelas et al., 2014). Proponents of this equilibrial paradigm concede that a precise mechanism explaining community regulation remains elusive (Magurran et al., 2018). Our numerical metacommunity model hints at a potential resolution to this problem and highlights an important avenue for the development of novel analytic theory. Inspection of recent results by Abernethy et al. (2019) for a spatially explicit food-web model in the light of our observations suggests that the phenomena we observe are not restricted to competitive communities, but may apply to a wider range of ecological models.

With this study we set out to assess the degree to which spatially unresolved ecological theory can incorporate the complex spatial processes occurring within model metacommunities. To our surprise, metacommunity models which explicitly incorporate dynamics at both local and regional scales reproduce an unprecedented range of empirically ubiquitous macroecological patterns. Crucially, these patterns result indirectly from the local dynamics, and in the neighbourhood of regional diversity limits. From this observation we conclude that there is an important interaction between the system-scale dynamical process central to the theory of ecological structural stability, and these key macroecological configurations. The spatial decoupling of timescales we observe (Fig 4A) implies that for a metacommunity of sufficiently large spatial extent regional ecological turnover could occur at time scales comparable to evolutionary or long-term environmental processes (e.g. glaciation cycles), as discussed by Ricklefs (2004). Because these processes are not include in our model, the ‘regional’ scale we refer to here must be understood as an intermediate spatial scale at which ecological processes operate comparatively fast.

If we conclude, on the basis of this and similar studies, that diversity regulation is indeed a common or general feature of ecological communities, this would entail a paradigm shift with important implications for the conservation and management of biodiversity. The assumption that local ecological dynamics have negligible impact on regional biotic distributions is still implicit in the majority of current conservation policies and programs. Species distribution modelling (SDM) is a widely used method for identifying ecological processes and responses of species distributions to environmental change. The basic SDM methodology assumes a comprehensive understanding of current and future climate is sufficient to predict range shifts under climate change. Our results suggest that ignoring biotic interactions, even if they cannot be explicitly detected using conventional tools, may strongly undermine the effectiveness of these models (Wisz et al., 2013). As such, we suggest that the development and application of more mechanistic distribution modelling (Dormann et al., 2018) should be a priority, that management models might focus on higher levels of biological organisation (e.g. feeding guilds or entire communities), and that designers of conservation and management strategies make a concerted effort to integrate factors relating to diversity regulation in their decision making.

## Methods

### Model landscape

We generated a spatial network consisting of *N* patches by sampling the Cartesian coordinates (*P*_*x*_, *Q*_*x*_) of each patch *x* (with 1 *≤ x ≤ N*) from a uniform distribution in the range 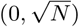. The local communities, were thus randomly distributed with density *≈* 1 over a model landscape of area *N*. Corridors were defined using the Gabriel algorithm (1969) which connects nodes *x* and *y* if the disc with diameter given by the line segment 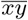 contains no other nodes. This non-trivial topography is more realistic than a complete graph, but was also selected for its relative computational efficiency. In the limit of large *N*, the average patch degree does not exceed 4 (Matula and Sokal, 1980), as for a square lattice, which will permit implementation of parallel simulation methods currently under development. Numerical experiments with fully connected graph lead to qualitatively similar emergent properties (see Supporting Information).

Environmental heterogeneity was modelled indirectly through spatial variation in species’ linear growth rates *r*_*ix*_, where the subscript *i* is a species index, and *x* a patch index. The values of *r*_*ix*_ were sampled from a Gaussian Random Field (Adler, 1981) (*μ* = 1.0, *σ*^2^ = 0.5), generated via spectral decomposition of the *N* by *N* landscape covariance matrix with elements **Σ**_*L,xy*_ = exp [*-ϕ*^-1^*d*_*xy*_], where *d*_*xy*_ denotes the Euclidean distances between two patches *x* and *y*, and the parameter *ϕ* controls the spatial autocorrelation of the environment (Johnson and Wichern, 2002). By varying the correlation length *ϕ* while keeping mean and variance of the fields fixed, we modelled landscapes of varying degrees of environmental heterogeneity (see Supporting Information). Parameters were chosen as *N* = 10, *ϕ* = 1 for Fig. 1, *N* = 20, *ϕ* = 1 for Figs. 4 and 5. (Temporal decoupling of spatial scales is clearer in larger regional communities, however for *γ*-diversity *≫ α*-diversity it becomes difficult to represent local and regional assemblages on a single axis.) Figs. 2, 3, and 6 summarize the complete parameter space studied: *N* = 2, 4, 6, 8, 10, 12, 15, 20, and 25; *ϕ* = 1, 2, 5, 10, 20, 40, 80, and 160, in all combinations. Landscapes of *ϕ>* 160 (in the spatial range studied here) showed no further decrease in *gamma*, or the exponent of the SAR, implying an effectively uniform environment.

### Dynamic equations and metacommunity assembly

We used a spatial extension of the Lotka-Volterra multi-species competition equation to model local population dynamics and dispersal in our model metacommunities, thus building on the model family pioneered by Reichenbach et al. (2007). The rate of change of local biomass of species *i* at patch *x* is given by the non-linear ordinary differential equation

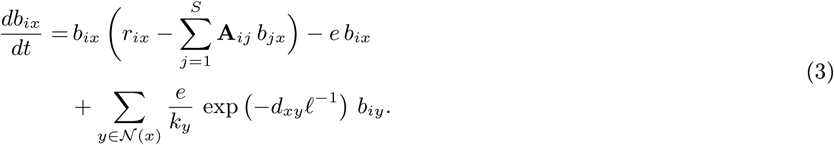

The system of *N × S* coupled equations can therefore be written as

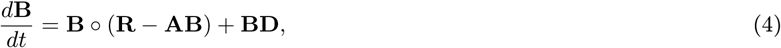

with ○ denoting element-wise multiplication.

The first term on the right hand side of Eq. (3) represents the local dynamics, where **A**_*ij*_ are the entries of the spatially unresolved competitive overlap matrix. In simulations, the off-diagonal entries **A**_*ij*_ were sampled randomly, with **A**_*ij*_ set to 0.3 with probability 0.3 and to 0 otherwise. The diagonal entries, representing intraspecific competition, were always set to 1. Together, Eq. (2) and Fig. 2 imply that the critical diversity at which a model community converges depends only on the expectation and variance of the regional interspecific interaction coefficients. The details of the distribution from which the **A**_*ij*_ are sampled do not enter the spatially implicit theory. For interaction matrices with pronounced structure (e.g. food webs), however, the underlying theory breaks down Rossberg (2013). The impact of the fundamental distribution from which the local interactions **A**_*ij*_ are sampled on spatial diversity patterns is subject of ongoing research.

The second term on the right of Eq. (3) represents the rate at which biomass of species *i* emigrates away from patch *x*, while the third term gives the immigration rates from all patches *y* sharing an edge with *x*. The immigration rate decays exponentially with characteristic length *ℓ*, kept fixed at 0.2. The parameter *e*, which represents the fraction of biomass leaving patch *x* per unit time, was kept fixed at 0.02. An exploration of the parameter space of *e* and *ℓ* revealed little qualitative shift in the emergent properties of the model over the biologically relevant range (see Supporting Information), thus values were selected that favoured computational efficiency during model assembly. The normalization constant *k*_*y*_ divides the biomass departing patches *y* between all other patches in the *its* local neighbourhood (𝒩 (*y*)), weighted by the ease of reaching each patch i.e. *k*_*y*_ =Σ_*z∈𝒩* (*y*)_ exp (-*d*_*yz*_*ℓ*^-1^)

We adopted the community assembly modelling approach first developed by Post and Pimm (1983). In each iteration of the algorithm, a new species was added to the metacommunity. Invaders were selected by computing the linear growth rate at low abundance of new species *i* with randomly generated ecologies (*r*_*ix*_ and **A**_*ij*_), until a species with positive linear growth rate in at least one patch was found. This was then added to the patch in which its effective growth rate was greatest with a low invasion biomass of 0.01 times the detection threshold of 10^-4^ biomass units. During invader testing the competitive impact of the invader on the dynamics of resident species was set to zero, such that resident biomass was unaffected, to make sure we capture the invader’s linear dynamics at low abundance. The metacommunity dynamics, including the spread of the new invader though the network and associated restructuring the local resident biomass distribution, were then simulated using the SUNDIALS numerical ODE solver (Hindmarsh et al., 2005) over 500 unit times, *t*. Those species whose biomass dropped below the detection threshold in all patches of the network were considered regionally extinct and removed from the system. By thus iteratively adding species to the community we modelled a constant flux of invaders, which causes the regional assemblage to self-organize, eventually converging on an equilibrium at which the invasion and extinction rates are equal on average. To reach this equilibrium, total simulation time was chosen as 4000, 6000, 8000, 10000, and 12000 iterations for *N ≤* 4, 6 *≤ N ≤* 10, 12 *≤ N ≤* 15, *N* = 20 and *N* = 25, respectively.

### Source-Sink classification

Source populations in a given patch *x* are those capable of locally maintaining themselves. Mathematically, source populations were defined as those for which local biomass was greater than the detection threshold and 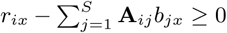. Conversely, sink populations where those of biomass greater than the detection threshold and 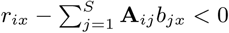.

### Regional scale interaction matrices, C

In order to compare model metacommunity dynamics to theoretical predictions, we numerically computed a spatially unresolved competitive overlap matrix, denoted **C**, that summarized the macroscopic dynamics at the regional scale. For this we constructed a spatially *unresolved* Lotka-Volterra system,

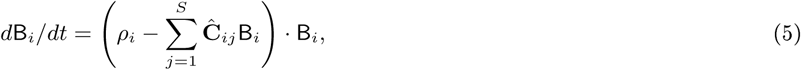

where B_*i*_ represents total biomass of species *i*, as an approximation of the spatially resolved model (Eq. (3)). The aim of the following method is to arrive at a description of the effective interaction between pairs of species given the self-organized spatial structure of metacommunity, which permits regional coexistence via spatial niche segregation. This requires integrating ecological interactions over the entire landscape, which was done using the computational equivalent of a harvesting experiment, under the assumption that interaction strengths can be inferred from the changes in regional abundances that result from controlled changes in the regional abundances of harvested species (Gilbert et al., 2014). Specifically, we asked how the steady state community responds to spatially unselective, light harvesting of a single species in the full model, and determined the coefficients **Ĉ**_*ij*_ of the unresolved model such as to obtain identical responses to linear order in the harvesting rate.

For the unresolved model, Eq. (5), the harvesting of species *i* at a rate *h*, represented by substituting *ρ*_*i*_ *- h* for *ρ*_*i*_ in Eq. (5), produces a shift in the equilibrium biomasses given by

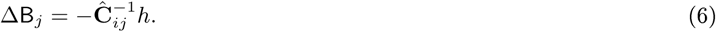

The most computationally efficient way of conducting the corresponding experiment for the meta-community is to use a numerical approximation of the Jacobian matrix. In doing so, we assume simulated metacommunities to be at fixed points, an approximation that is justified retrospectively by the apparent efficacy of the method. The elements of the Jacobian are given by the general equation

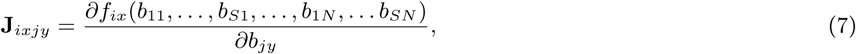

evaluated at equilibrium. The functions *f*_*ix*_ denote the right hand side of Eq. (3).

Light harvesting of a single focal species *i* at a rate *h* brings about a small shift in the equilibrium biomasses of the other species in the metacommunity and the dynamics of the harvested community *near the unharvested equilibrium* 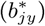 can be approximated by

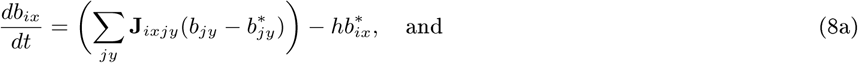

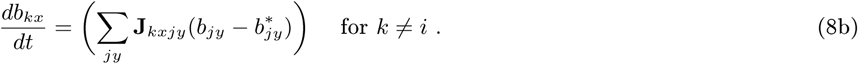

Here *h* is the harvesting rate. We vectorize the matrix **B** (denoted 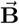) such as to match the dimensionality of the spatially resolved Jacobian, and the write the equilibrium condition for Eq. (8a) as

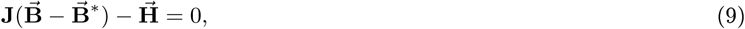

where the elements of vector 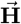 are 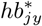 for *j* = *i* and 0 otherwise. From this, we obtain

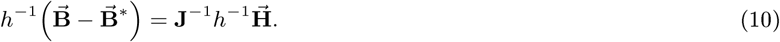

The left hand side of Eq. (10) represents the local shift in biomasses due to the harvesting of the focal species *i* per unit *h*. From Eq. (10) we compute the change in total biomass 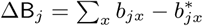 for each species *j*. Comparison with (6) gives row *i* of **Ĉ** ^-1^. Iterating over all species *i* = 1 *…S*, we computed **Ĉ** ^-1^ and from this the spatially unresolved interaction matrix **Ĉ**. Finally, in order to match the assumptions made in the derivation of Eq. (2) (Rossberg, 2013), we divided each row and column **Ĉ** by the square root of the corresponding diagonal element to obtain the *effective competitive overlap matrix*, **C** (which has ones along the diagonal).

### Temporal diversity patterns

Temporal species richness and turnover in community composition were computed for a metacommunity at regional diversity limits for a period corresponding to 500 ecological invasions (Fig. 4). Species richness analysis requires the application of some presence-absence criterion. We assess local community diversity by reference to the source populations only, since sink populations are effectively decoupled from local filtering processes by dispersal.

Following the 500 invasions, species were removed in reverse order of regional abundance, in order to model a large scale mass extinction process. A single metacommunity of *N* = 20, *ϕ* = 2 was used for this analysis.

Compositional turnover in metacommunities at regional diversity equilibria was measured using the Bray-Curtis (1957) similarity. An arbitrary initial metacommunity composition was selected (*T* = 0 in Fig. 4A) and the relative compositional change computed in the context of a constant invasion flux using the function **vegdist** in the R package “vegan” (Oksanen et al., 2018). In order to generate Fig. 4A, regionally excluded or as yet uninvaded species were assigned biomass vectors with all elements set to zero.

### Species ranges

In order to quantify range sizes of species, we first computed, for each species, the population covariance matrix

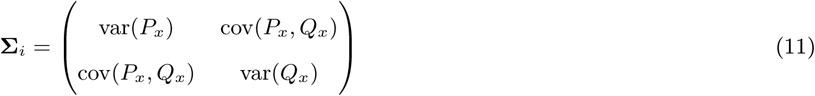

of the locations (*P*_*x*_, *Q*_*x*_) of individuals forming the species’ population, assuming population sizes are proportional to biomasses at each patch. For example, with *i* being the index of the focal species, 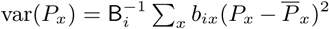, where B_*i*_ = Σ_*x*_ *b*_*ix*_ is total biomass, as above, and 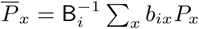 is the *P* component of the centre of mass of the distribution. Correspondingly,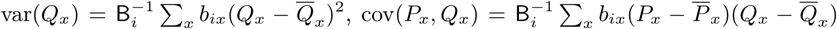, with 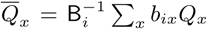. As a measure of range size, we computed the product of the square roots of the eigenvalues of **Σ**_*i*_, i.e. the square root of its determinant,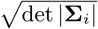. By this measure, an even distribution of biomass over the full 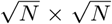 rectangle enclosing one of our model communities corresponds to a range size of *N/*12.

### Species co-occurrence

In order to analyse the correlation in species spatial distributions within our model landscapes, we used the probabilistic model developed by Veech (2013) included in the R package “cooccur” (Griffith et al., 2016). The observed pair-wise co-occurrence is computed as the probability of detecting species *i* in patch *x* given the detection of species *j* in that local community, which is then compared to that expected if two species were distributed independently within a discretized landscape.

