## Supporting Information for "Metacommunity-scale biodiversity regulation and the self-organized emergence of macroecological patterns"

### The coupling of local and regional competition coefficients

The interaction between environmental gradients and competitive overlap coefficients in the determination of the regional overlap coefficients $\mathbf{C}_{ij}$ is complex and depends upon the degree of environmental heterogeneity. To test the hypothesis that local interactions ($\mathbf{A}_{ij}$) drive regional community assembly (Rabosky and Hurlbert, 2015) we need to analyse the extent to which fundamental and realised competitive overlap coefficients are correlated. Simple linear regression shows that $\mathbf{A}_{ij}$ and $\mathbf{C}_{ij}$ are indeed significantly correlated ($p < 0.01$, for all parameter combinations), but the strength of the correlation, measured by the adjusted $R^2$, ranges from 0.1 to 1.0.

Unsurprisingly, given the nature of $\mathbf{C}$, the slope of the relationship $m \leq 1$, with $m \sim 1$ in the homogeneous extreme (high $\phi$ and/or low $N$, Fig. S1A), in which case the strength of the correlation between local interactions and regional community assembly is maximised (Fig. S1B).

### Spatial structure at the metapopulation scale

The combined effect of local and regional diversity regulation drives spatial turnover in community composition. Species in model metacommunities near regional limits converge on highly aggregated spatial distributions in which biomass decays exponentially as a function of distance from regional maxima (Fig. S2, note the logarithmic colour gradient).

### Regional scale competitive overlap matrices

Fig. S3A-B highlights the difference in magnitude of the regional scale interaction coefficients, which incorporate the complex spatial coexistence mechanisms active in heterogeneous landscapes. Spatial coexistence mechanisms in model metacommuities can be understood as slowing the onset of structural instability by modifying the distribution in spatially resolved competitive overlap coefficients. The spectrum of the 'microcosmic' competitive overlap matrix $\mathbf{A}$ (Fig. S3C), fully covering the origin of the complex plane, highlights the fact that such an assemblage would be biologically unfeasible in the absence of the abiotic (topographical and environmental) variation in the model.

### Spatial autocorrelation in maximum intrinsic growth rates

Spatially correlated intrinsic growth rates $r_{ix}$ were modelled using a Gaussian random field as described in *Model landscape*. Fig. S4 shows four complete realizations of spatially distributed growth rates, $r_{ix}$, generated using autocorrelation lengths $\phi$ =1 and 10.

### Dispersal parameterization and landscape topography

The numerical experiments conducted in this study, focused on a single point in the parameter space of $e$, and $\ell$ (0.02,0.2). In practicse the parameterization was selected in order to maximize the computational efficiency of the assembly process. Here we show that the self-organized spatial structure of strongly regulated model metacommunities is robust to changes in the dispersal parameters within a biologically meaningful parameter space. Figs. S5 and S6 show the regional diversity, average local source and sink richness, and the skew of the RSD, for a model metacommunities ($N = 20$, $\phi = 1$) assembled using parameters $e$ and $\ell$ spanning four orders of magnitude, with each parameter combination triple replicated.

In both cases the regional diversity increased slightly as a function of $\ell$ in the case $e < 1$. This results contrasts with that observed, for example, by Mouquet and Loreau (2003), who find that in the high dispersal regime, metacommunities tend to converge on monoculture. In our model system, environmental heterogeneity (here high), ensures that no species can dominate the entire landscape. The weak *increase* in diversity suggests that dispersal limited rare species may be rescued from regional extirpation by increasing landscape connectivity. In the case $e > 1$, which implies $e > \bar{r}_{ix}$, the regional diversity is significantly reduced and an approximate monoculture does indeed emerge in the high dispersal regime, as the metacommunity approaches the patch-dynamics paradigm assumed by Mouquet and Loreau (2003).

The average local source richness is largely unaffected by dispersal parameterization in the case $e < 1$ since this component is instead strongly regulated by local ecological dynamics. In the case $e > 1$, all populations depend in immigration from adjacent patches and the average local source diversity drops immediately to zero.

The average local sink richness depends strongly on parameterization, and landscape topography. For $e < 1$ sink richness increases as a function of dispersal length until the combined source and sink diversity approaches on the regional species richness, a scenario in which most species have a detectable population in all local communities. This result is qualitatively comparable to that observed by Bastolla et al. (2001), who showed how island diversity increases as a result of super-saturation due to fast immigration from a continental species pool. In the case $e > 1$ it is difficult to meaningfully interpret source-sink dynamics, since in this extreme dispersal regime, all populations depend on immigration for persistence. It is only in this extreme case that we observe consistent deviation from what is arguably the key macroecological property of our model metacommunities - a regional biota with an RSD that is positively skewed.

Our study focuses on the parameter space in which $\gamma$-diversity is large compared to mean $\alpha$-diversity, implying meaningful spatial turnover in community composition. While investigation of systems in which dispersal effectively homogenizes local communities is interesting, it is outside of the scope of this study. In the parameter space of interest here (the highlighted region in Figs. S5 and S6) we consider the diversity patterns to be qualitatively robust.

The choice to employ the Gabriel graph *as well as* an exponential dispersal kernel, enforces dispersal limitation in two potentially conflated ways. As described in section *Model landscape*, this was largely a pragmatic choice. We are confident, however, that macroscopic properties of the metacommunity do not depend on the topography of the network. Comparison of Figs. S5 and S6), shows how, the non-trivial patch adjacency reduces average sink diversity by effectively reducing the average range within the assembly. The qualitative patterns, however, are not strongly impacted and as such we surmise the emergent phenomena depend more on the ecological, than the topographic network.

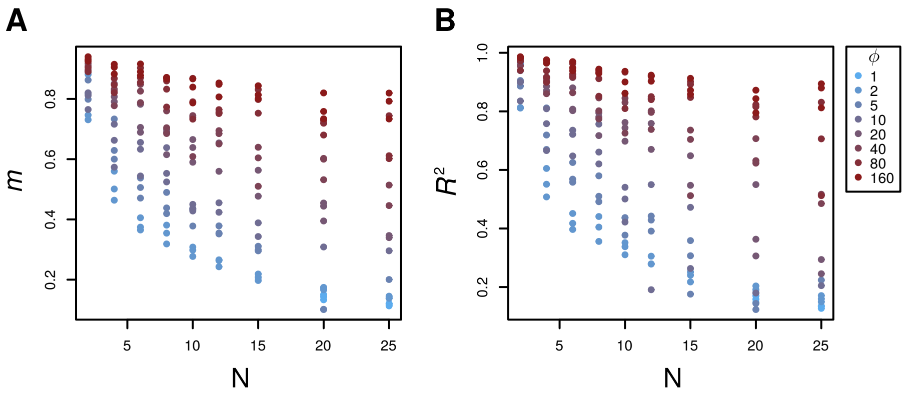


Figure S1: **The statistical relationship between interspecific competitive overlap coefficients $\mathbf{A}_{ij}$ and $\mathbf{C}_{ij}$.** The slope of the linear regression of the elements of $\mathbf{C}_{ij}$ against those of $\mathbf{A}_{ij}$ (A) and the adjusted $R^2$ of the relationship (B). Colours denote to environmental auto-correlation length, $\phi$.

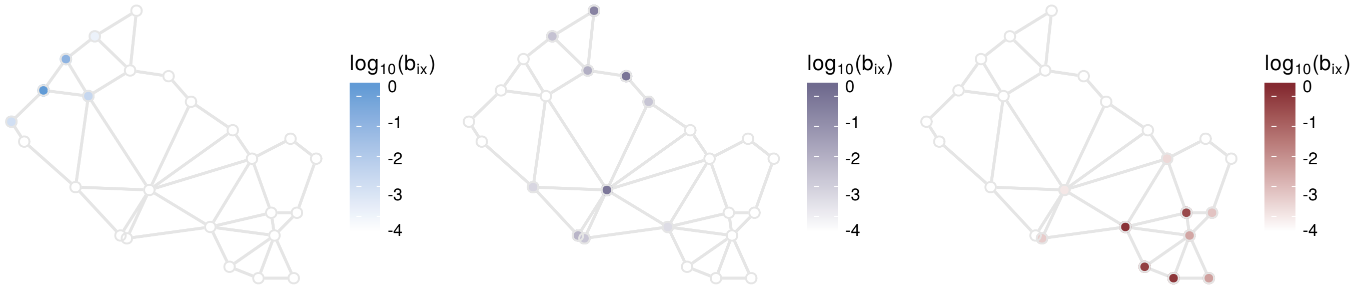


Figure S2: **A typical spatial model landscape and the distribution of biomass for three simulated species.** Dispersal corridors (gray links) in random spatial networks were assigning using the Gabriel Algorithm. Colours correspond to three randomly selected species biomass distributions. Spatial aggregation is a key characteristic of species ranges in strongly regulated metacommunities.

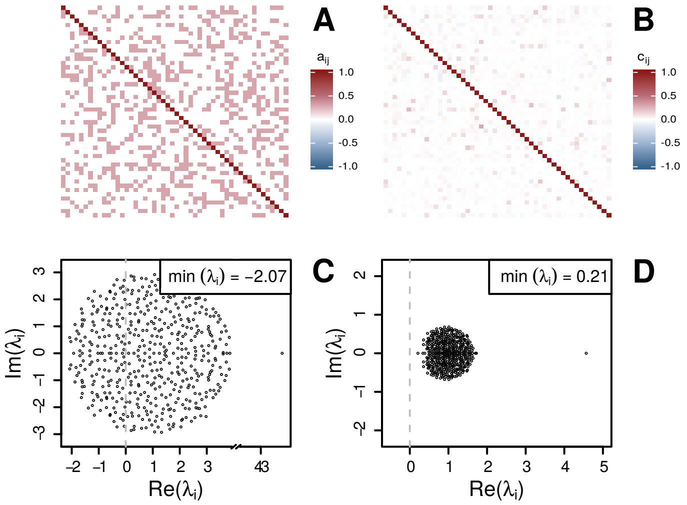


Figure S3: **Comparison of the matrices A and C for a single metacommunity at regional diversity equilibrium.** The local competitive overlap matrix **A** (A) and regional *effective* competitive overlap matrix **C** (B) for a randomly sampled subset (for visual clarity) of a typical model metacommunity ($N = 20$, $\phi = 1$) at regional diversity equilibrium and the corresponding eigenvalue spectra (C and D). The off-diagonal elements, $\mathbf{C}_{ij}$, incorporate the effects of both spatial segregation and 'indirect mutualism' (which can lead to negative values). The spectrum of the regional overlap matrix **C**, which approaches the origin as diversity saturates, matches that observed in spatially unresolved models (Rossberg, 2013).

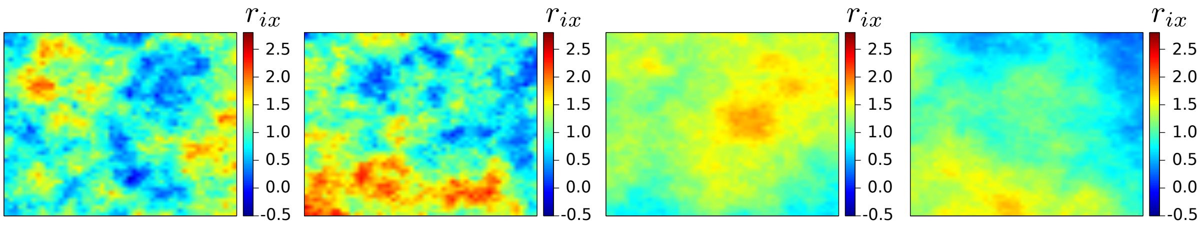


Figure S4: **Spatially autocorrelated Gaussian random fields used in environmental modelling** Four complete realization of Gaussian random fields generated via eigen-decomposition of the spatial covariance matrix. The 'landscapes' shown here have side lengths of $\sqrt{20}$ and spatial correlation lengths, $\phi$, of 1 (left) and 10 (right).

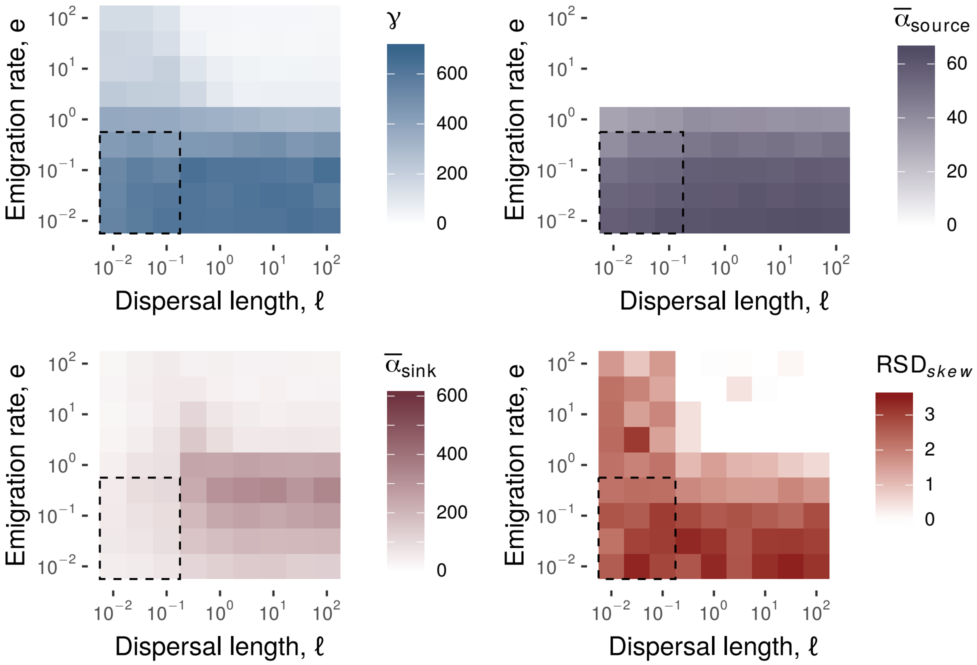


Figure S5: **Regional, average source, and sink species richness, and the skewness of the RSD under variation of** $e$ **and** $\ell$**, for the Gabriel graph**. Exploration of the parameter space of the dispersal parameters shows that our results are qualitatively robust to variation in $e$ and $\ell$ in the biologically relevant regime indicated by the dashed box. See text for further details.

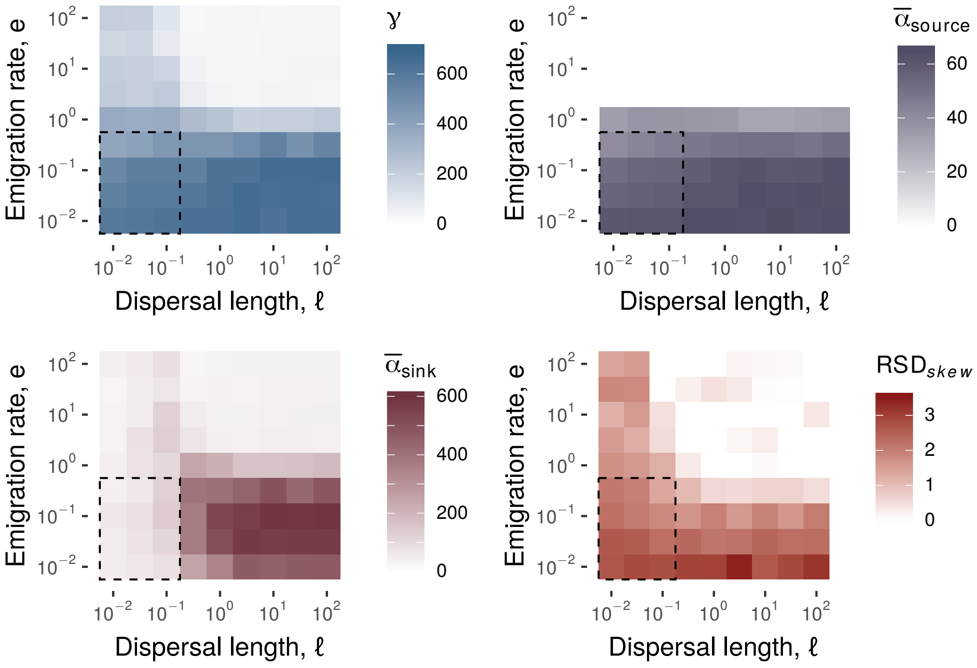


Figure S6: **Regional, average source, and sink species richness, and the skewness of the RSD under variation of $e$ and $\ell$, for the Complete graph**. As in Fig. S5, for fully connected random graphs.
